# Not every gene is special: one simple rule to control the false discovery rate when analysing high-throughput sequencing data

**DOI:** 10.1101/2025.11.28.690516

**Authors:** Scott J. Dos Santos, Andreea C. Murariu, Justin D. Silverman, Gregory B. Gloor

**Author notes:** Equal contribution (Scott J. Dos Santos, Andreea C. Murariu).

## Abstract

Differential expression and differential abundance analyses are commonplace in studies employing high-throughput sequencing approaches; however different tools often fail to return comparable results when applied to the same dataset. Most tools employ various normalisations to attempt to correct for technical variation in the count data. Previously, we demonstrated that these normalisations are often inappropriate due to incorrect assumptions regarding the overall scale (i.e. size) of the biological system in question. In this study, we conducted 100 permutation analyses of 10 RNA-seq datasets to show that scale misspecification results in poor control of the false discovery rate by several commonly-used analysis tools. Moreover, we demonstrate that this can be ameliorated by using a scale model in ALDEx2 or ALDEx3 to account for uncertainty around the size of the system scale. We show that there is an inherent trade-off between satisfactory control of false-discovery rates and sensitivity and that no tool offers both. We also quantify how increasing scale uncertainty affects the difference between groups required for a feature to be reported as differentially expressed. Finally, we provide universal guidance on choosing an appropriate amount of scale uncertainty for any type of analysis. Overall, our work highlights the strengths and pitfalls of commonly used tools for differential expression analyses and highlights the choice between sensitivity and false-discovery rate control that all researchers are making when analysing sequencing data.

## 2 Introduction

High-throughput sequencing (HTS) has been widely adopted across the biological sciences in the study of health and disease [1, 2]. Various approaches exist, including amplicon sequencing using universal ‘barcoding’ genes (e.g. 16S rRNA gene hypervariable regions [3], *cpn*60 [4], *rpoB* [5]), shotgun metagenomic sequencing [6], bulk and single-cell RNA-seq [7, 8], and metatranscriptome sequencing [9]. Determining the taxa, genes or functions that distinguish two different biological conditions of interest is a common end-point for such analyses and a multitude of tools have been developed to enable this [10, 11].

Tools for differential expression/abundance analysis typically take as input a table of read counts per gene or taxon across all samples. In reality, this count table does not represent the absolute abundance of the given features in their original environment [12, 13].The counts represent relative abundances on account of several steps in the sequencing workflow being inherently compositional [14]. Only a minute proportion of the final, pooled sequencing library is loaded onto the sequencing flow cell which itself originates from a small amount of each original DNA extract, itself a fraction of the original sample and so on. All information regarding the true biological scale of the community under study is lost and all downstream analyses are ignorant to this information, regardless of the tool used [15]. Ultimately, variation in sequencing depth across samples from the same dataset is decoupled from and unrelated to system scale.

Most analyses involve some sort of normalisation of the read count table in the workflow to account for differences in sequencing depth; however, this unwittingly introduces a critical problem [16]. Normalisations often make an inaccurate assumption about system scale [17, 18, 19]. Examples of such assumptions include: i) that the geometric mean of the observed data accurately reflects system scale [20]; ii) that the total microbial abundance is the same across all samples [21]; iii) that certain features within a sample represent a suitable reference that may be used to scale others [22, 23]; iv) that scale remains constant across different parts of a sample [24]. Recent work has shown that violating these assumptions can inflate the false discovery rate (FDR) beyond what can be considered reasonable; in fact the true FDR can be above 75% in some cases [25, 26, 18, 19].

Poor control of the FDR is a widespread issue across many experimental designs that use HTS data; indeed, the problem of poor FDR control in high dimensional biological data dates to at least the analysis of microarray data [27]. Poor FDR control is a long-standing problem when analysing bulk transcriptome data [28, 25] and similar issues arise with microbiome data [29, 30]. In response to this, the use of a dual cutoff approach where both a *P*-value and a log_2_ fold-change (L2FC) are commonly used to help reduce the number of positive identifications [27]. Specific guidance for transcriptome data was provided by Schurch *et al*. using a large bulk transcriptome dataset [31]. This dual cutoff method is widespread in the HTS literature, and its adoption was enhanced by the very useful volcano plot [27] that is now standard across many fields that use HTS.

However, soon after the dual cutoff approach was first used, it was shown not to control the FDR [32]. There are several reasons for this. First, there is a mismatch between the assumptions of variance for the two approaches; the L2FC cutoff assumes that all genes have the same variance, while the statistical test uses a gene-specific variance to determine the *P*-value. One way to think about this is that there are two ways that a gene can be significantly different between groups but undesirably so; it can have a low variance and small L2FC, or it can have a large variance and large L2FC [33]. Second, while the widely-used Benjamini-Hochberg [34] (BH) procedure controls the FDR for all rejected genes, it does not necessarily do so over any subset of genes, and indeed false positives are often over-represented in any such subset [35, 33]. One way to deal with the first problem is shrinkage to provide better variance estimates [32], and shrinkage is implemented in popular tools such as DESeq2 [36], and limma [37]. However, there is no way to deal with the second issue, as is apparent by the inflated FDR exhibited by many, if not all tools as noted above.

To address the important issue of poor FDR control, Nixon *et al.* [17] proposed an alternative to normalisation-based methods, introducing the concept of scale models through a framework known as ‘scale reliant inference’ (SRI). These scale models have been incorporated into the ALDEx2 R package [19], a Bayesian toolbox for modelling the underlying variation in sequencing count data and estimating the log_2_ fold change (L2FC). Originally, ALDEx2 applied a centred log-ratio (CLR) transformation to many instances of the modelled count data drawn from a Dirichlet distribution [38]; however in doing so, it made the implicit, yet incorrect assumption that system scale is equivalent to the inverse of the geometric mean used for calculating the CLR (see work by Nixon *et al*. [17] and McGovern *et al*. [39] for details). Following the incorporation of scale models into ALDEx2, system scale is now directly modelled across each separate Dirichlet instance, acknowledging the reality that a single point-estimate of scale is essentially guaranteed to be incorrect [19]. By accounting for scale uncertainty, this modification converts the original ALDEx2 algorithm into a model termed a ‘scale simulation random variable’ [17]. ALDEx3 is a new implementation of the ALDEx2 approach; ALDEx3 is based on linear models and requires fewer computational resources while still implementing the Bayesian approach to modelling and analysis.

Konnaris and colleagues demonstrated the utility of scale models by applying ALDEx3 to one of the largest collections of paired microbiome-microbial abundance data to date [26], comparing its performance to machine learning (ML) models from a recent high-profile study [40]. These ML models failed to generalise to new datasets from the *mutt* database [26] when predicting microbial load and also failed to improve on the FDR achieved by scale-naïve ALDEx3. In contrast, ALDEx3 scale models achieved a superior FDR, and better positive- and negative-predictive values when applied to both shotgun metagenomic data and 16S rRNA amplicon sequencing data. Similar findings have been reported for ALDEx2 when applied to a variety of HTS datasets; superior control of the FDR is achieved when applying the SRI framework [19].

Previous work highlighting the advantages of the SRI framework has mainly focused on amplicon sequencing and shotgun metagenomic datasets [26], or has been limited to a single transcriptomic dataset [19]. Here, we extend prior studies by testing the performance of scale models from ALDEx2, and its faster and more memory-efficient successor, ALDEx3, on 10 different RNA-seq datasets, encompassing approaches such as bulk transcriptome (from both lab-derived and clinical samples), single-cell and metatranscriptome sequencing. We compare these two implementations of SRI to widely-used, normalisation-based tools in terms of sensitivity and FDR, and highlight an important trade-off between the two that researchers should consider when analysing HTS data. Further, we quantify the relationship between the degree of scale uncertainty modelled by ALDEx2 and ALDEx3 and the minimum fold-change between conditions required for a statistically significant result-a critical factor to consider when conducting differential expression analysis with scale models. This analysis provides strong guidance on how to build appropriate scale models that offer better FDR controls for HTS data analysis.

## 3 Materials and methods

### 3.1 Datasets used in analysis

We obtained read count data and grouping metadata from publicly available RNA-seq datasets described by the following studies or sources: Li *et al.* [25] (PD1 immunotherapy dataset), Gierlinski *et al.* [41] (wild-type vs. *Snf2*-knockout yeast transcriptome dataset), Skinnider *et al.* [42] (single-cell RNA-seq dataset comparing cytotoxic T cells vs. memory T cells), and Dos Santos *et al.* [43] (vaginal metatranscriptome dataset comparing healthy vs. bacterial vaginosis patients, aggregated by KEGG orthology terms). A further seven bulk RNA-seq datasets originating from the Cancer Genome Atlas project [44] were obtained from the supplementary data provided by Li *et al*. via Zenodo [45] (available at: https://zenodo.org/records/8320659), corresponding to the following tumour types: breast invasive carcinoma (BRCA) [46, 47], kidney renal cell carcinoma (KIRC) [48], liver hepatocellular carcinoma (LIHC) [49], lung adenocarcinoma (LUAD) [50], prostate adenocarcinoma (PRAD) [51], and thyroid carcinoma (THCA) [52].

All Cancer Genome Atlas datasets, the PD1 immunotherapy dataset, and the yeast transcriptome dataset were filtered using the filterByExpr() function from the edgeR package [53]. The single-cell RNA-seq dataset was filtered such that only 1000 randomly selected cells in the upper quartile of read counts were retained in each group, along with features with read count totals of at least 300 reads across all samples. The vaginal metatranscriptome dataset was filtered using the CoDaSeq package (v0.99.7) [54] to retain features present in 30 % of samples, at a minimum proportion of 0.005 % of a given sample (as per the original study) [43].

### 3.2 Binomial thinning of count data

For the analysis of all datasets, we applied binomial thinning via the seqgendiff R package (v1.2.4) [55] to achieve the following: i) permute the grouping variable such that group membership of all samples is randomised; and ii) inject an artificial signal into read count data such that 5 % of all features in a given dataset are modelled to be differentially abundant between groups. The result of the former is that any analysis with the permuted grouping variable is meaningless as almost all positive findings are, by definition, false-positives (FPs; excluding features affected by binomial thinning). The result of the latter is that any features whose counts are altered by binomial thinning *could* be detected as being significantly different between groups and are considered simulated ‘true’ positives (TPs). The magnitude of the difference betwee groups (i.e. the amount of ‘signal’ added) is drawn from a normal distribution with a standard deviation of 2 and a mean of zero; because of this, not all features with a modelled difference are actually detectable (i.e. false negatives, FNs). Features whose counts have not been altered by binomial thinning and are not identified as significantly different between groups are ‘true’ negatives (TNs).

### 3.3 Calculation of sensitivity and false discovery rate

For each iteration of the permutation analyses, sensitivity (i.e. true positive rate, TPR) was calculated as the number of TPs reported by a given tool as differentially expressed, divided by the total number of simulated TP features altered by binomial thinning in a given analysis iteration which equates to:

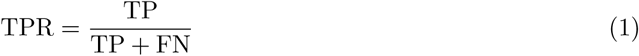

Likewise, the false discovery rate (FDR) was calculated for each analysis iteration as the number of features incorrectly defined by a tool as differentially expressed (i.e. reported as significantly different but not among the features altered by binomial thinning), divided by the total number of differentially expressed features reported by that tool, equating to:

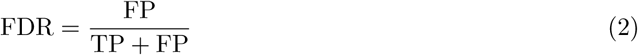

At each 0.1 increment from 0 to 1, the features which were altered by binomial thinning must have a simulated (i.e. ‘true’) log_2_ fold-change exceeding this value to be considered a TP; this threshold is referred to as the model difference between groups. For every model difference threshold from 0 to 1, sensitivity and FDR were calculated as above for each tool and dataset, based on the number of TP features for that threshold. For example: for a feature to be considered a TP at a model difference threshold of 0.5, the log_2_ fold-change ‘injected’ by binomial thinning must be ≥0.5, and for the same feature to be considered a positive finding by a given tool, that tool must report a corresponding Benjamini-Hochberg (BH) adjusted *P*-value ≤0.05 [34]. Note that BH-adjusted *P*-values are calculated only once per analysis iteration for the entire set of features. Positive predictive values were also calculated for each tool and dataset as 1 − FDR.

### 3.4 Differential expression analyses

We tested the performance of ALDEx2 (v1.4.1) [38], ALDEx3 (v0.3.0) [26], DESeq2 (v1.49.1) [36], edgeR (v4.7.2), and limma (v3.65.1; limma-voom) [37] across the aforementioned 10 datasets in RStudio, using R version 4.5.1 [56]. A total of 100 group permutations were performed for each dataset and tool combination. For each analysis iteration, group membership permutation and binomial thinning were performed after setting a dynamic seed value to ensure unique, yet reproducible, permutations (seed value = 20 + iteration number). Analysis functions were run on permuted, thinned data using the permuted grouping variable after setting a static seed value to ensure reproducibility.

### 3.5 Modelling scale uncertainty with ALDEx2 and ALDEx3

The ALDEx2 and ALDEx3 packages model uncertainty in the scale of the biological system under study, as has been extensively detailed elsewhere [18, 19]. Briefly, we can describe the true counts of any system, **W**, which contains *D* features and *N* samples, as the product of the proportional parts of the samples (i.e composition) and the total counts of the samples (i.e. the scale). Thus, any single sample from system **W** can be described fully by Eq. 3.

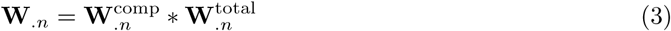

Unfortunately, we can never measure the true system directly, owing to the compositional nature of HTS data which provide only an indirect readout of the compositional information of, system **W** [15, 57]. The observations we produce by sequencing for a given feature in a single sample is described as **Y***_dn_*. Each observation is an integer representation of the probability of measuring a given feature in the given sample, scaled by the total number of reads collected[58]. The single point observations from **Y***_dn_* are converted to a Bayesian posterior distribution of proportions [58] to provide a distribution of reasonable estimates. This is done by sampling from a Dirichlet distribution to give a posterior estimate of 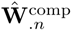 [38] (where the ^indicates an estimate) as in Eq. 4:

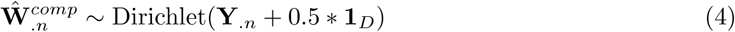

Data from HTS are always normalised to remove nuisance parameters and make them commensurate [22]. The insight of Nixon et al. [17] was that almost all normalisations made the strict assumption of being a direct estimate of the sample system scale: that is, that 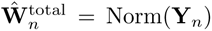. Nixon and colleagues showed that these estimates were always incorrect, but could be made more useful by including uncertainty. The Bayesian framework was adapted to include scale uncertainty in the normalization as in Eq 5.

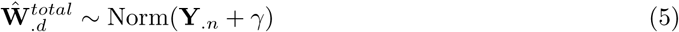

Here, *γ* is drawn from a Gaussian distribution with mean *µ_n_* and variance 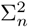. In this way the full posterior estimate of each sample in **Ŵ** incorporates uncertainty in both the observed data and the normalisation of that data as in Eq. 6.

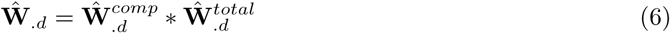

Programmatically, *γ* is controlled by the ‘`gamma`’ parameter in the call to ‘`aldex.clr()`’. This parameter can be used to incorporate only uncertainty in the estimation of normalisation, or can be tuned to include prior information about absolute abundance in each sample along with the error of those observations. See Nixon et al. [18], for a fulsome description of the `gamma` parameter. For both ALDEx2 and ALDEx3, simple scale models can be constructed by passing a single numeric value to the ‘`gamma`’ argument. These scale values will be used to adjust the log-ratio transformed count data. In the present work, gamma values from essentially zero (0.001) to 0.5 in increments of 0.1 were used for both tools to model biological system scale.

### 3.6 Data availability and visualisation

All code needed to reproduce these analyses are available on GitHub at: https://github.com/amurariu/usri. This repository also contains input count data and metadata for all datasets, as well as code needed to replicate the figures in this study. All figures were produced using ggplot2 (v3.5.2) [59] and edited in Inkscape (v1.3.1). For transparency, details of figure edits made in Inkscape are available on GitHub, in addition to the original SVG files.

## 4 Results

### 4.1 High false discovery rates are universal across scale-naïve tools

Since there is usually no objective standard of truth for high throughput data, we calculated the false discovery rate (FDR) for all combinations of tools and gamma values using a strategy of group permutation with the injection of artificial ‘true’ signal into 5% of all features using binomial thinning [55]. In every dataset tested, DESeq2, edgeR and limma-voom exhibited the highest FDRs of all tools, compared to scale-naïve and scale-aware ALDEx2 and ALDEx3 (Fig. **1**). DESeq2 and edgeR routinely returned FDRs that were often double or more the reported nominal values, and were always larger than the FDR reported by either ALDEx2 and ALDEx3. This problem was most acute in Cancer Genome Atlas datasets (Supplementary Fig. **S1**) and was not as pronounced in the yeast transcriptome dataset. Both ALDEx2 and ALDEx3 demonstrated better FDR control in all datasets, even when system scale was not modelled. Increasing scale uncertainty further reduced the FDR, such that when γ ≥0.1 there were virtually no false discoveries until the modelled log_2_ fold-change (L2FC) between groups reached approximately 0.5; this was universal across all datasets. At higher modelled difference thresholds, the effect of scale was even more pronounced when compared to scale-naïve analyses: the FDR was below 5% (or practically zero, in many cases) when γ ≥0.4, regardless of the L2FC simulated by binomial thinning. Of the three tools unable to incorporate scale information, the lowest FDR was generally seen with limma; the sole exception being for the single-cell RNA-seq dataset. Universally, the FDR increased as the L2FC threshold concomitantly increased, with FDRs being highest when considering features with the largest L2FCs between groups. FDRs routinely exceeded 20 %, and could exceed 40 %, when features had a modelled L2FC *>*1. This was particularly noteworthy given that a large fold-change between groups is often interpreted as a sign that expression of a particular feature is both truly different between conditions and also important to the biological situation under study. Naturally, the same patterns are seen when the positive predictive value is calculated (Supplementary Fig. **S2**).

**Figure 1:**
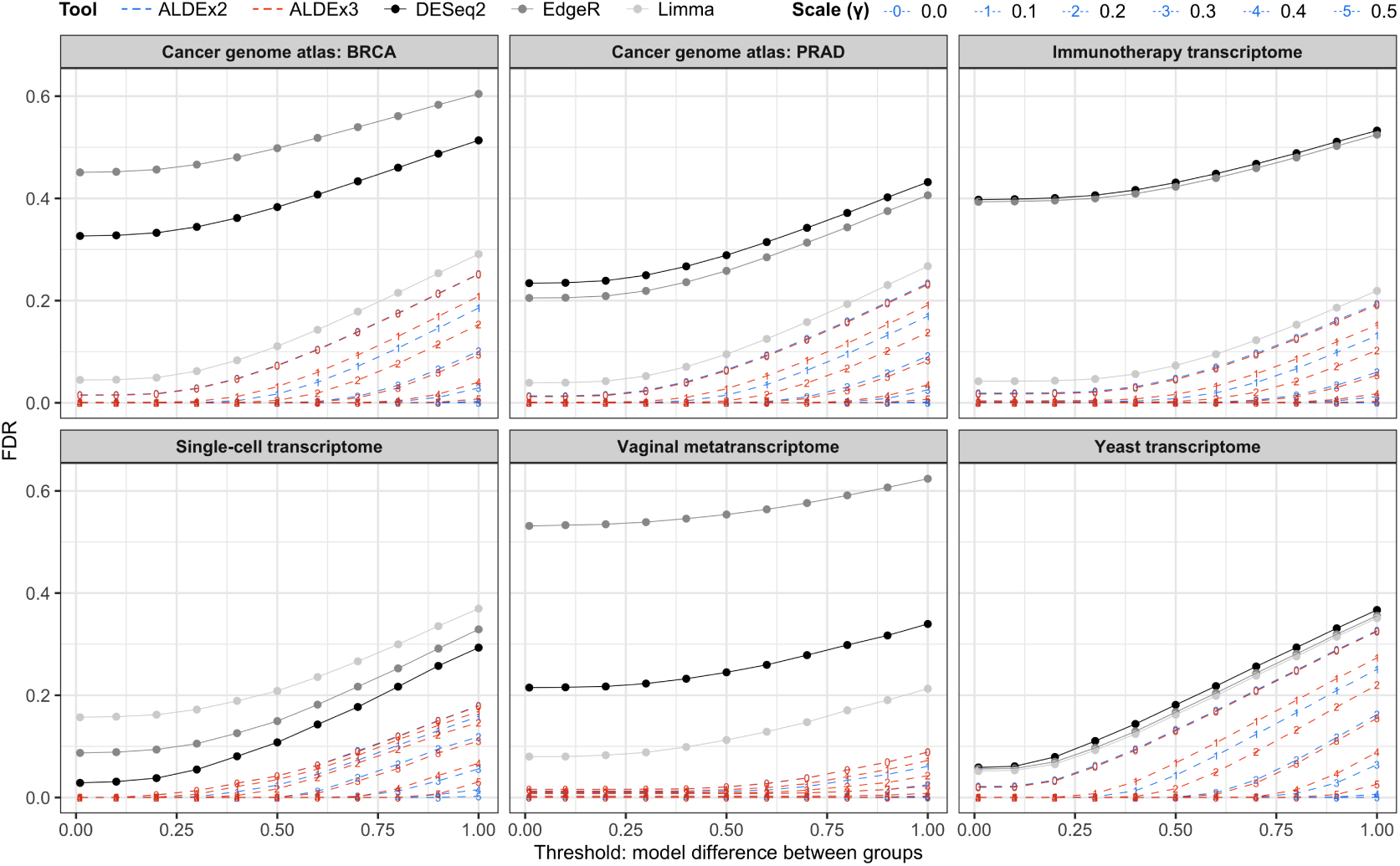
Scale-naïve analyses provide poor control of the false discovery rate (FDR) across all datasets. False-discovery rates for the indicated tool and dataset combinations were calculated separately for each 0.1 increment in the model difference between groups threshold (i.e. the simulated or ‘true’ log_2_ fold-change between groups, as determined by binomial thinning). Each data point represents the FDR when considering dataset features with a simulated log-difference at or above the given x-axis threshold.

### 4.2 Commonly used tools offer a trade-off between high sensitivity and good FDR control

We next calculated the true-positive rate (TPR), or sensitivity, for all tool and dataset combinations. Unlike what was observed for the FDR calculations, DESeq2, edgeR and limma exhibited higher sensitivity in every dataset tested than did either version of ALDEx (Fig. **2**). Conversely sensitivity was lower for both ALDEx2 and ALDEx3 across every dataset, even in scale-naïve analyses. This was expected because the ALDEx model is a consensus model [38] and generally has a somewhat lower sensitivity [29, 30]. As was observed for FDR control, the addition of scale uncertainty had an evident effect on TPRs: increasing gamma values reduced sensitivity, with larger values (γ ≥0.4) resulting in less than 50 % of true positive features be-ing detected by ALDEx2 in some datasets. ALDEx3 performed better than its predecessor at every degree of scale uncertainty tested, remaining competitive with DESeq2, edgeR and limma when small amounts of scale were added (γ ≤0.2). Notably, ALDEx2 and ALDEx3 had poor sensitivity with the single-cell dataset and to a lesser extent, the vaginal metatranscriptome dataset (though the latter may be due to the fact that a simple scale model is insufficient for the vaginal metatranscriptome; see discussion). TPRs for these datasets were markedly lower than the remaining bulk transcriptome data (Supplementary Fig. **S1**), failing to achieve *>*40 and *>*50 %, respectively, in many cases. Notably, the effect of the modelled L2FC on sensitivity was again obvious. The TPR tended to plateau for all tools when the modeled L2FC was ≥0.5, although at high γ values ≥0.4, this was not the case.

**Figure 2:**
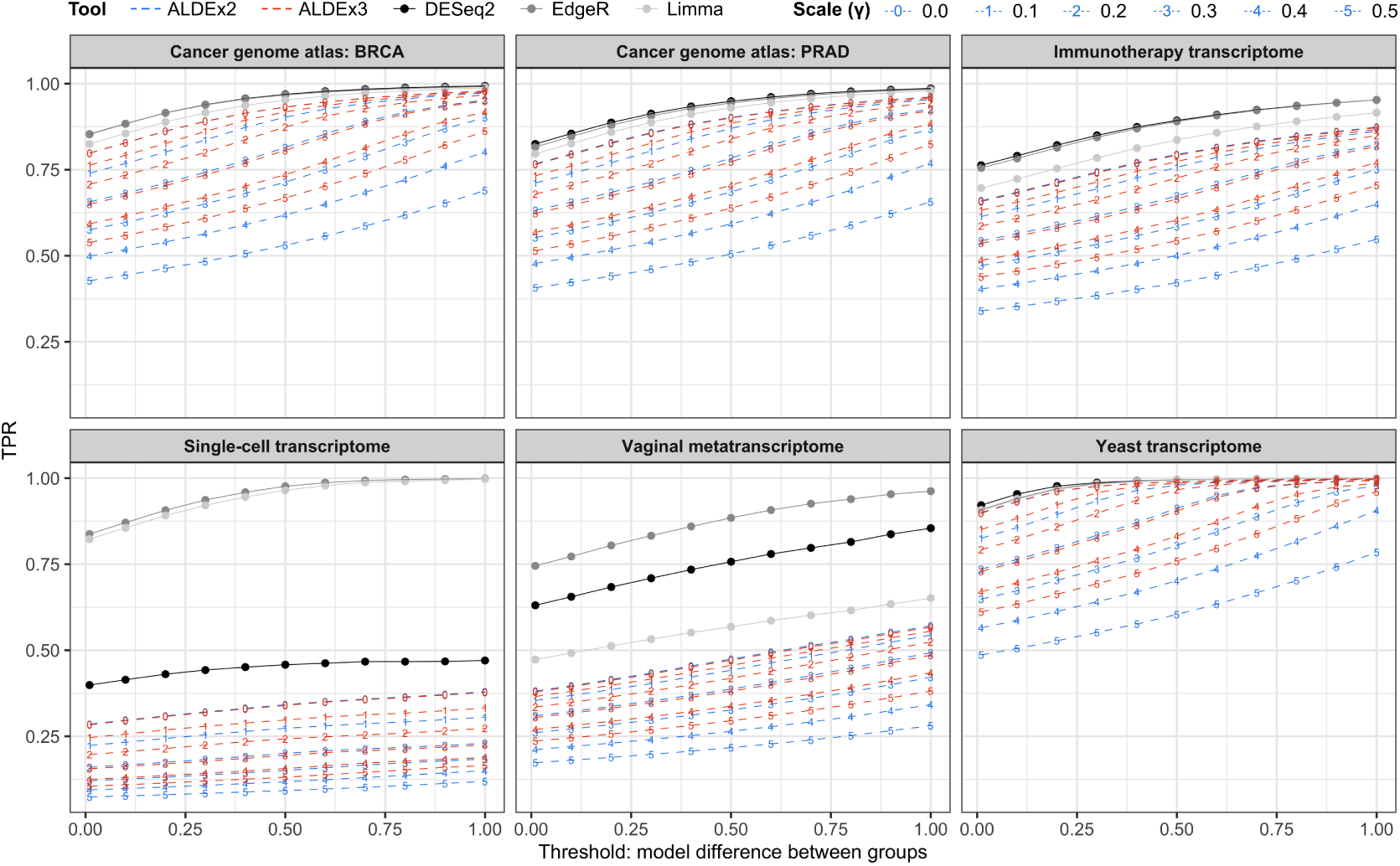
Sensitivity is reduced when adding increasing amounts of scale uncertainty. True positive rates (TPR; i.e. sensitivity) for the indicated tool and dataset combinations were calculated separately for each 0.1 increment in the model difference between groups threshold (i.e. the simulated or ‘true’ log_2_ fold-change between groups, as determined by binomial thinning). Each data point represents the TPR when considering dataset features with a simulated log-difference at or above the given x-axis threshold.

### 4.3 Increasing scale uncertainty increases the minimum fold-change required for significance

Using the PD1 immunotherapy dataset as an example, we investigated the effect of increasing values of γ in ALDEx2 and ALDEx3 relative to the scale-naïve analysis (i.e., with γ set to 0.001). The reduction in sensitivity from increasing γ was accompanied by a reduction in the total number of differentially expressed features returned for both tools (Table **1**). ALDEx2 was more sensitive to this effect than was ALDEx3, mirroring the results of the sensitivity and FDR benchmarking.

**Table 1:** Number of differentially expressed genes reported by ALDEx2 and ALDEx3 for the immunotherapy dataset. Data expressed as the mean of all 100 iterations rounded to the nearest integer, with standard deviations given in parentheses.

| Scale uncertainty ($\gamma$) | No. significant genes (± s.d.) | |
| --- | --- | --- |
|  | ALDEx2 | ALDEx3 |
| 0 | 689 (60) | 685 (56) |
| 0.1 | 628 (20) | 647 (21) |
| 0.2 | 554 (22) | 599 (20) |
| 0.3 | 481 (24) | 547 (22) |
| 0.4 | 412 (26) | 497 (22) |
| 0.5 | 347 (29) | 447 (22) |

To explain both the decrease in significantly different features and the superior FDR control that characterise ALDEx2 and ALDEx3, we constructed volcano plots [27] (difference between groups vs. −log_10_ *p*-value) and effect plots [60] (difference within groups vs. difference between groups) from a randomly chosen analysis iteration of the PD1 immunotherapy dataset. For both tools, the volcano plots show distinct, curved boundaries between genes that were significantly different between groups at increasing amounts of scale uncertainty (Fig. **3**A and **3**B). The same approximate boundaries for each γ value in additional datasets can also be seen in Supplementary Fig. **S5**. By explicitly modeling uncertainty in the scale, both tools eliminated potentially spurious false-positive results by a dynamic cutoff that: i) required a large L2FC if near the FDR cutoff value chosen (i.e. features have a large L2FC but marginal statistical significance) or, ii) allowed a smaller L2FC value if the adjusted *p*-value was smaller (i.e. features are more statistically significant, but not necessarily biologically pertinent). The same phenomenon is illustrated in effect plots (Fig. **3**C and **3**D): features that exhibit a larger dispersion within a group require a larger difference between groups to remain significantly different at larger γ values. This effect is less pronounced with ALDEx3, explaining its increased sensitivity even at higher degrees of scale uncertainty.

**Figure 3:**
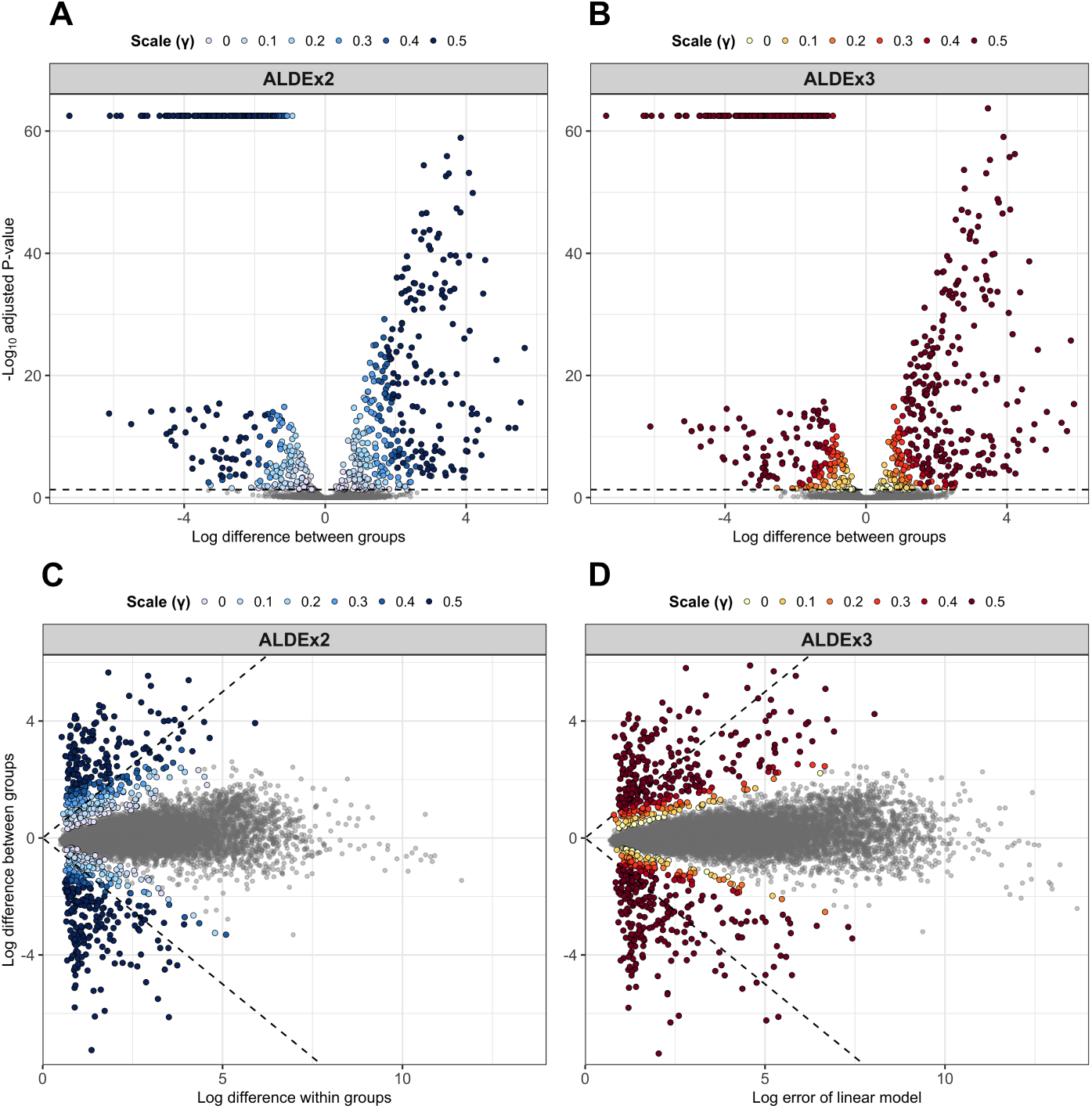
Accounting for scale uncertainty reduces the total number of features identified as differentially expressed. A randomly selected iteration from the ALDEx2 and ALDEx3 PD1 immunotherapy dataset analyses was used to create volcano plots (**A, B**) and effect plots (**C, D**). Each data point represents a feature from the scale-naïve analysis (γ = 0.001). Grey points indicate non-significant features; coloured points indicate features which are significantly different between groups after a given amount of scale uncertainty has been added to the model. Grey dashed lines indicate either: Benjamini-Hochberg-corrected *p*-value *<*0.05 (**A, B**) or equivalent difference between and within groups (**C, D**).

To further understand the effect of increasing scale uncertainty on differential expression analyses, we visualised the minimum L2FC required for ALDEx2 and ALDEx3 to return a statistically significant result across all 100 iterations of the PD1 immunotherapy dataset (Fig. **4**A). At almost every γ value, the mean L2FC required for a significant result was lower with ALDEx3 compared to ALDEx2– the sole exception to this being the scale-naïve analyses, where the difference was negligible (0.34 ALDEx2 vs. 0.36 ALDEx3; Supplementary Table **S1**). The distribution of minimum differences needed for a significant result became markedly wider as the degree of scale uncertainty increased with ALDEx2. However, for ALDEx3 these distributions were comparatively narrower for all values of γ tested.

**Figure 4:**
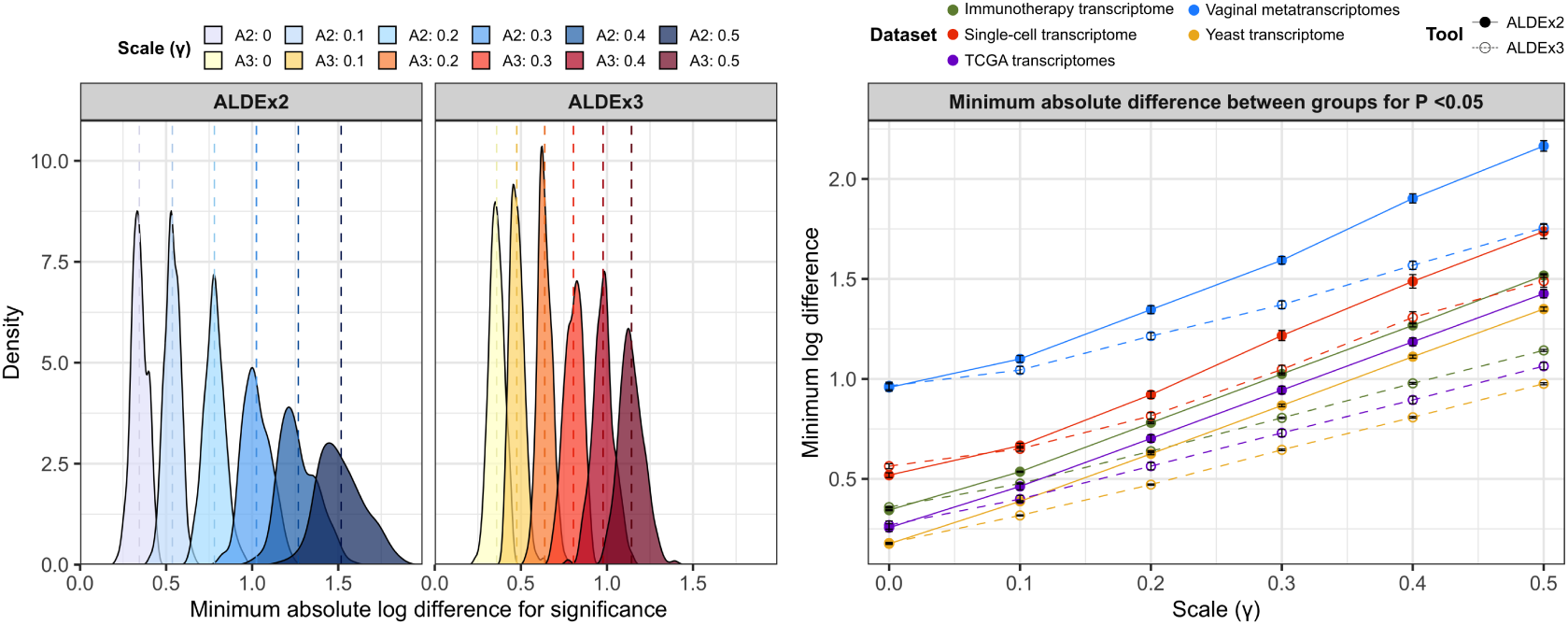
ALDEx3 requires a lower difference between groups to identify differentially expressed genes compared to ALDEx2. (**Left**) For each of the 100 iterations of the PD1 immunotherapy transcriptome dataset, the minimum absolute difference between groups needed for a gene to be reported as differentially expressed by ALDEx2 and ALDEx3 was recorded, based on a BH-adjusted *P*-value of *<*0.05, for γ values ranging from 0 to 0.5. The distributions of these values are coloured by tool and γ value, with dashed lines indicating the mean. (**Right**) Minimum absolute differences required for a significant result were calculated across all 100 iterations for all datasets. Mean values per dataset are plotted (± SEM) for ALDEx2 and ALDEx3. Cancer Genome Atlas datasets are aggregated and plotted data represent a mean of means at each γ value.

We next examined the minimum L2FC difference as a function of the amount of scale uncertainty added across all datasets to determine the generality of the observations in the PD1 immunotherapy dataset (Fig. **4**B). We observed that different datasets had different L2FC thresholds for statistical significance even in the absence of additional scale uncertainty. For all datasets, the additional L2FC required for statistical significance was again most noticeable at the largest degree of scale uncertainty tested. The L2FC threshold for significance was associated with the amount of measurement variation in the datasets: those that have very little measurement uncertainty (yeast, TCGA) having small initial thresholds and those with substantial uncertainty (single-cell, metatranscriptome) having larger initial thresholds (Supplementary Fig. **S3**). Strikingly, for both ALDEx2 and ALDEx3 we observed the relationship between the minimum L2FC and the additional scale uncertainty to be nearly constant for each tool, although the slopes differed considerably by tool. The distribution of the minimum difference required for significance was narrower for bulk RNA-seq datasets analysed with ALDEx2 compared to metatranscriptomic or single-cell data; however, all distributions were comparatively narrower when data were analysed with ALDEx3 (Supplementary Fig. **S4**). Metatranscriptomic data in particular exhibited a relatively large amount of variability in the minimum log difference required for a significant result between iterations. Linear regression of these average minimum differences required for significance vs. gamma values of 0 to 0.5 similar slopes for all tool and dataset combinations, averaging to a mean of 2.39 for ALDEx2 and 1.64 for ALDEx3 (Fig. **4**B).

Table **2** shows slopes and intercepts for the minimum L2FC when scale uncertainty is added to the various datasets. This was calculated as the mean coefficient and intercept from linear regression models of the minimum log differences required for significance vs. γ values of 0 to 0.5, across all 100 iterations of ALDEx2 and ALDEx3 analyses, for all datasets. As in Fig. **4**B, slopes for ALDEx2 ranged from 2.34 to 2.53, while slopes for ALDEx3 ranged from 1.60 to 1.95. Thus, we concluded that there was a near-constant relationship between increasing scale uncertainty and the additional L2FC required for significance across disparate data types. The utility of this relationship in selecting an appropriate value of γ is expanded upon in the discussion.

**Table 2:** Linear models of gamma values vs. minimum log differences required for significance over 100 iterations of ALDEx2 and ALDEx3: The minimum log difference required for a statistically significant result (*P <*0.05) was calculated for each iteration of the ALDEx2 and ALDEx3 analyses for all datasets. Linear regression was performed on these minimum differences vs. the corresponding gamma values and the mean coefficients and intercepts of the models are reported. Values in parentheses represent standard deviations.

| Dataset | ALDEx2 (± s.d.) | | ALDEx3 (± s.d.) | |
| --- | --- | --- | --- | --- |
|  | Coefficient | Intercept | Coefficient | Intercept |
| Immunotherapy | **2.37** (0.28) | **0.32** (0.03) | **1.60** (0.15) | **0.33** (0.03) |
| Metatranscriptome | **2.49** (0.90) | **0.89** (0.23) | **1.62** (0.82) | **0.91** (0.25) |
| TCGA: BRCA | **2.37** (0.28) | **0.20** (0.03) | **1.61** (0.15) | **0.21** (0.02) |
| TCGA: KIRC | **2.35** (0.28) | **0.20** (0.03) | **1.62** (0.14) | **0.21** (0.03) |
| TCGA: LIHC | **2.39** (0.29) | **0.29** (0.04) | **1.61** (0.16) | **0.31** (0.04) |
| TCGA: LUAD | **2.35** (0.27) | **0.24** (0.03) | **1.61** (0.16) | **0.26** (0.03) |
| TCGA: PRAD | **2.34** (0.28) | **0.21** (0.03) | **1.61** (0.16) | **0.22** (0.03) |
| TCGA: THCA | **2.35** (0.27) | **0.21** (0.03) | **1.61** (0.14) | **0.22** (0.03) |
| Single-cell | **2.53** (1.11) | **0.46** (0.16) | **1.95** (0.97) | **0.49** (0.19) |
| Yeast | **2.37** (0.30) | **0.16** (0.03) | **1.61** (0.15) | **0.16** (0.03) |

### 4.4 Applying scale models to real datasets highlights spurious findings in scale-naïve analyses

Having shown the advantages and trade-offs of modelling system scale in semi-simulated datasets with binomial thinning, we next applied scale models to the original, unpermuted BRCA dataset to assess the performance of ALDEx2 and ALDEx3 on real data. In a scale-naïve analysis of healthy vs. tumour samples, ALDEx2 identified 15,837 differentially expressed genes, whereas ALDEx3 identified 15,825. For comparison, analyses with DESeq2, edgeR and limma all yielded a larger number of differentially expressed genes (16,736, 16,407 and 16,295 respectively). As expected, increasing amounts of scale uncertainty reduced the number of differentially expressed genes reported by both ALDEx2 and ALDEx3, down to 2,777 and 4,437 respectively at γ = 0.5 (Supplementary Table **S2** & Supplementary Fig. **S5**).

Critically, for each increase in scale uncertainty, the overwhelming majority of genes dropped by ALDEx2 and ALDEx3 (*>*99 %) did not belong to the set of experimentally validated breast cancer drivers [61]. Those that did tended to have a low BH-adjusted *P*-value, a low fold-change between groups, a high within-group dispersion, or a combination thereof (Supplementary File S1). For example, expression of the well-known tumour suppressor gene, *PTEN*, was no longer significantly different between groups according to ALDEx2 at a γ value of 0.1. Notably, the log_2_ fold-change was minute (−0.0594), the BH-adjusted *P*-value was already marginal in the scale-naïve analysis (*P* = 0.034) and log within-group dispersion was relatively large (0.757). These observations suggested considerable heterogeneity in the expression of *PTEN* in this dataset. Examination of the raw and normalised expression values for *PTEN* (Supplementary Fig. **S6**) showed that the raw or normalised distributions overlapped nearly perfectly, with any perceived differences between tumor and normal adjacent tissue in this dataset being driven by a small number of outlier samples. Similar behaviour for this gene with the addition of scale uncertainty was seen with ALDEx3 (Supplementary File S1).

At higher degrees of scale-uncertainty, the reduced sensitivity of ALDEx2 became apparent. At γ = 0.5, genes known to be involved in cancer pathogenesis, such as *ERBB2* and *KRAS*, were no longer identified as differentially expressed, despite a two-fold difference in expression between groups. While RNA-seq data does not directly account for heterozygosity or non-functional proteins, these observations highlighted the problem of an excessively high γ value, and suggest that lower values are appropriate in most instances.

## 5 Discussion

The analysis of ‘-omics’ data is not, and likely will never be, standardised; each analyst has their preferred approach and tool, accompanied by its various pros and cons. This variation often leads to poor reproducibility, conflicting results, or both–even when the same dataset is analysed with different tools. One reason for this is that there has been no compelling reason to choose one approach over another, in part because of a lack of understanding as to why different tools give different answers. This has now changed with the introduction of the scale model approach [17] where it was shown that different normalisations made different assumptions about the sampled environment; these assumptions were shown to be always provably incorrect. The use of incorrect assumptions lead to false positive and false negative errors [17, 18, 19, 62].

Li *et al*. [25] used an approach similar to this work and found that both DESeq2 and edgeR often identified as many differentially abundant genes when analysing permuted datasets (i.e. all results are false-positives) as when analysing the original, unpermuted data. Results from each tool poorly overlapped with the other, particularly when considering the PD1 immunotherapy dataset used in the present work; only 8 % of differentially expressed genes were reported by both tools. Conversely, and worryingly, there was considerable overlap between the list of differentially expressed genes from analysis of permuted and original count data, highlighting the prevalence of false discoveries in any given DESeq2 or edgeR analysis. Among the false-positive findings were genes related to immune functions which, given the biological context, would seem like a plausible result; however in reality, none of the randomly permuted differences were true.

In this study we substantially extend our previous work across an additional 10 datasets of different data types, three additional tools, and 10-fold more permutations of the datasets. Here, we used a data permutation approach [25] to produce known false-positives, along with binomial thinning [55] to introduce potential true-positives. This combination allowed us to benchmark tools on data that exhibit all of the uncertainty and unpredictability inherent in real HTS data that is missing from pre-specified models, while still introducing a ‘true’ signal [55]. From this new data we identify universal guidance for the amount of scale uncertainty to add, and strengthen the argument that one cannot have one’s statistical cake and eat it. The sensitivity and FDR tradeoffs of some of the most widely used RNA-seq analysis tools are highly biased towards sensitivity, as has been observed before [29, 30, 25]. Thus, analysts can either be practically assured of getting a “statistically significant” result but at the risk of a very high chance of a false discovery, or they can expect lower sensitivity balanced by the confidence that any significant results represent a real difference between their biological conditions of interest.

In line with previous benchmarking on different types of datasets [29, 30, 25], we observed that both edgeR and DESeq2 often exhibit very high FDR rates that are balanced by very high sensitivity. We note that limma performs well in some but not all types of data, having an acceptable FDR and a high sensitivity in the majority of datasets. ALDEx2 and ALDEx3 have the lowest sensitivity overall, but also markedly lower FDR rates for all datasets. Notably, ALDEx3 provides the same functionality as its predecessor in terms of acknowledging biological scale during normalisation; however, it offers superior sensitivity at the same degree of scale uncertainty while also controlling for false discoveries at any given L2FC and γ value.

Both our work and that of Li *et al*. make the counterintuitive observation that the FDRs are highest when filtering for a larger L2FC in the permuted datasets. Prior work has shown that placing emphasis on such features through the common practice of double-filtering results for high fold-change and BH-adjusted *P*-values using volcano plots can inflate the FDR even further [27, 63]. These simulation experiments underscore the danger of taking the results of RNA-seq analysis tools at face values without considering the well-documented problem of inadequate FDR control [64, 32]. One obvious issue of using a simple-fold-change cutoff is that a feature can appear to have a large L2FC between groups, when in fact that change is driven by a few outlier samples. We therefore caution any user against associating large fold changes in expression with true biological differences simply because of the magnitude of difference without further inspection.

The incorporation of modeled scale uncertainty into the core ALDEx2 and ALDEx3 algorithms offers excellent FDR control with only modest loss of sensitivity. As noted, the use of a dual cutoff approach composed of *P*-value cutoffs and fold-change thresholds can inflate FDRs; however, even modest amounts of scale uncertainty eliminate the need for such an approach without inflating the FDR. The addition of scale uncertainty provided a way to exclude two types of features. First, those that have marginal BH-adjusted *P*-values but a large between-condition differences. Such features have large or unpredictable dispersions and often come from distributions where there are only a few outlier (or high leverage) samples represented. In our analysis of real-world data, the *PTEN* tumour suppressor gene was not differentially expressed when we added even a small amount of scale uncertainty; inspection of the raw input and normalised output confirmed it as an example of a gene where a few outlier samples drove the significant result. Second, many features have very small differences but minute BH-adjusted *P*-values. These represent features with a very small difference between groups and very small dispersion and so are likely not to be biologically relevant. Indeed, these are the same features that are excluded with a dual L2FC cutoff and *P*-value cutoff. Thus, the use of a scale model excludes both types of features that Ebrahimpoor et al. [33] suggests are problematic, without depending on subsetting the adjusted *P*-values, which is known to be counter productive [35, 33].

Although only simple scale models are explored in the present work, ALDEx2 and ALDEx3 are capable of constructing informed scale models that apply different degrees of scale uncertainty to each group of interest [17, 18]. We have previously shown that this approach is necessary in multiple vaginal metatranscriptome datasets to correct for the 10 to 100-fold difference in total bacterial load that exists between states of health and bacterial vaginosis [43, 65]. Using an informed scale model removes an asymmetry in the data that causes a multitude of housekeeping functions to be identified as differentially expressed, while also preventing known pathways involved in BV pathophysiology from being identified as such. Moreover, this methodology was shown to be robust to replication even when applying it to independent validation datasets with vastly different population demographics.

Notably, this work provides a path forward to reducing the FDR in a way that only minimally impacts sensitivity across many different types of HTS data. The relationship between γ and the minimum L2FC required for a statistically significant result shown in Fig. **4** and Table **2** provides almost universal guidance on selecting an appropriate level of scale uncertainty when using ALDEx2 or ALDEx3. We recommend that investigators introduce scale uncertainty into any analysis sufficient to provide a buffer of an additional 0.5-fold change. This equates to adding a γ value of 0.2 when using ALDEx2, and of 0.3 when using ALDEx3, and this ensures that the FDR rate is controlled at least to an inferred L2FC of *>*0.5 across all the datasets tested here with only minimal impacts on sensitivity. If the analyst is interested in only very large L2FC values *>*1, then we suggest using a γ value of 0.4 for either tool.

Overall, we have demonstrated the utility of scale models implemented by ALDEx2 and ALDEx3 in the analysis of high-throughput sequencing datasets, as well as their excellent FDR control. Like others, we highlight the unacceptably high rate of false-positive findings returned by commonly used RNA-seq analysis tools and note the inherent trade off all researchers need to be cognisant of when considering how to analyse their data.

## 6 Supplementary material

**Supplementary Figure S1:**
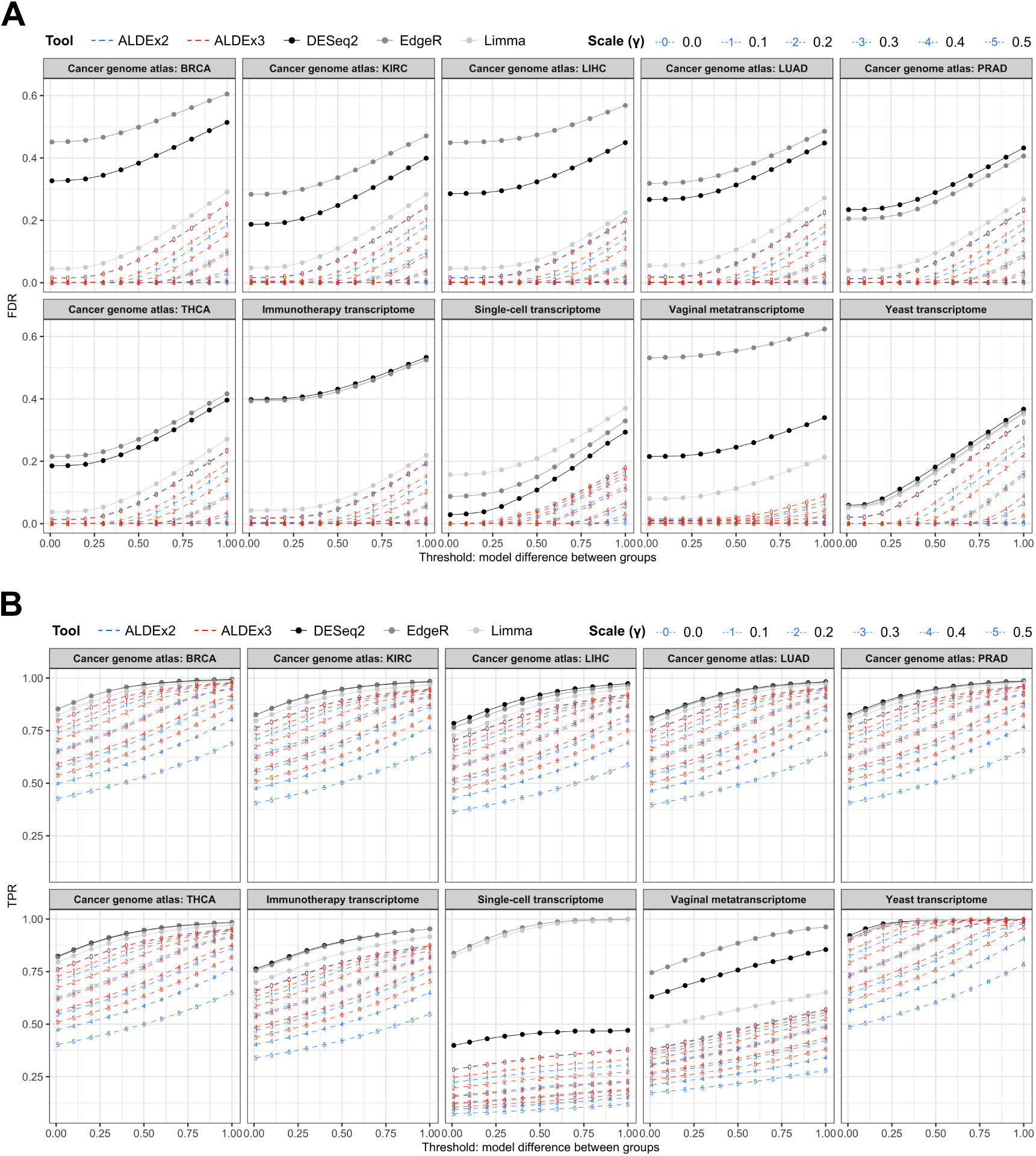
False-discovery and true-positive rates for all datasets tested: False-discovery rates (FDR; **A**) and true positive rates (TPR; **B**) for all tool and dataset combinations were calculated separately for each 0.1 increment in the model difference between groups (i.e. the simulated or ‘true’ log_2_ fold-change between groups, as determined by binomial thinning). Each data point represents the FDR or TPR when considering dataset features with a simulated log-difference at or above the given x-axis threshold.

**Supplementary Figure S2:**
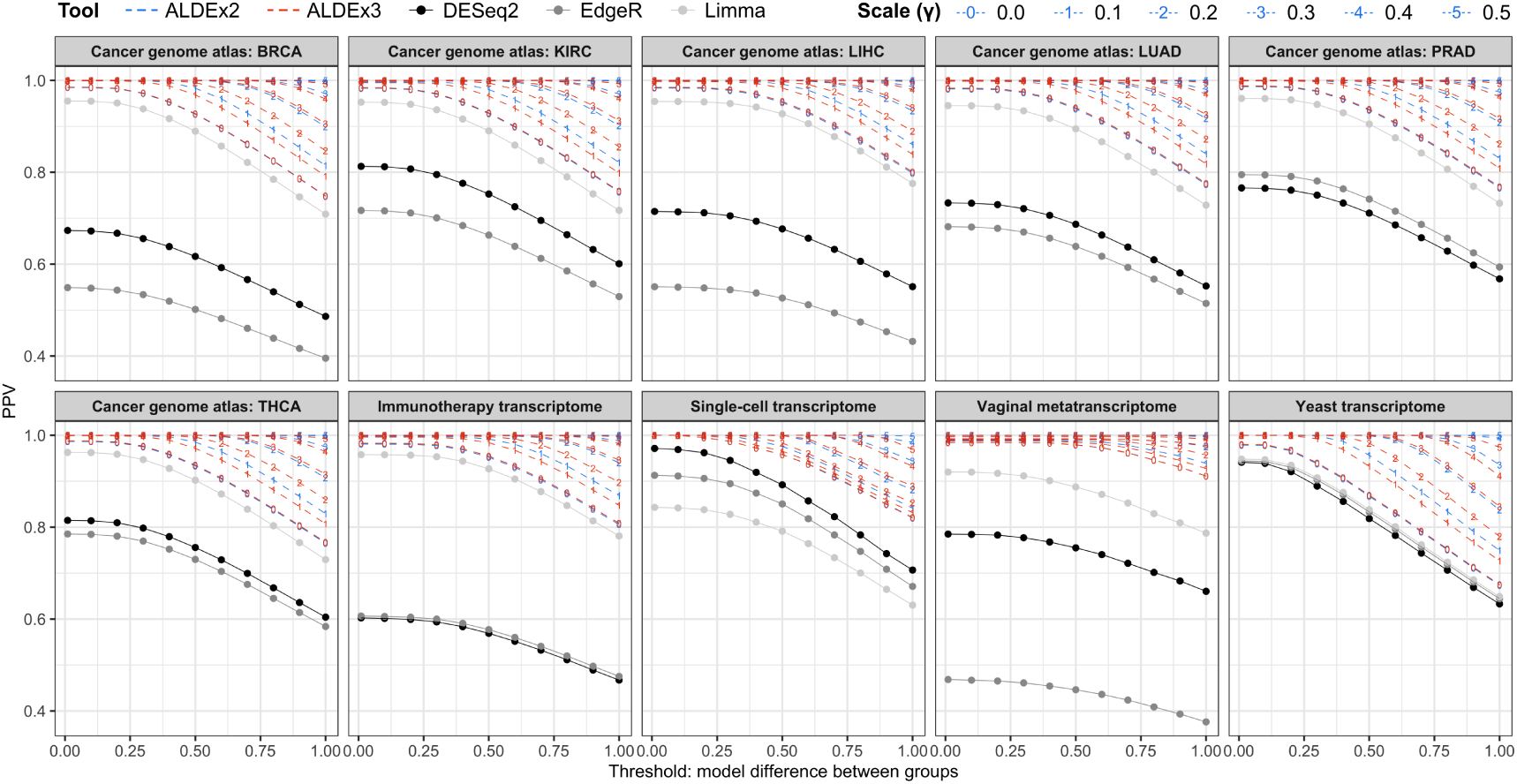
Positive predictive values for all datasets tested: The positive predictive value (PPV) for all tool and dataset combinations was calculated separately for each 0.1 increment in the model difference between groups (i.e. the simulated or ‘true’ log_2_ fold-change between groups, as determined by binomial thinning). Each data point represents the PPV when considering dataset features with a simulated log-difference at or above the given x-axis threshold.

**Supplementary Table S1:** Average minimum log differences required for a significant result with ALDEx2 and ALDEx3 for all datasets and γ values. For all datasets, the mean minimum log difference required for ALDEx2 and ALDEx3 to recognise a feature as differentially expressed was calculated for γ values ranging from 0 to 0.5, across all 100 iterations (TGCA, The Cancer Genome Atlas).

| Dataset | Scale ($\gamma$) | Mean minimum difference (± s.d.) | |
| --- | --- | --- | --- |
|  |  | ALDEx2 | ALDEx3 |
| Immunotherapy | 0 | **0.34** (0.045) | **0.36** (0.041) |
|  | 0.1 | **0.54** (0.044) | **0.48** (0.040) |
|  | 0.2 | **0.78** (0.059) | **0.64** (0.040) |
|  | 0.3 | **1.02** (0.079) | **0.80** (0.051) |
|  | 0.4 | **1.27** (0.108) | **0.98** (0.061) |
|  | 0.5 | **1.52** (0.132) | **1.14** (0.070) |
| Metatranscriptome | 0 | **0.96** (0.261) | **0.96** (0.260) |
|  | 0.1 | **1.10** (0.246) | **1.04** (0.257) |
|  | 0.2 | **1.35** (0.280) | **1.21** (0.245) |
|  | 0.3 | **1.59** (0.275) | **1.37** (0.271) |
|  | 0.4 | **1.90** (0.327) | **1.57** (0.291) |
|  | 0.5 | **2.16** (0.376) | **1.76** (0.291) |
| Single-cell | 0 | **0.52** (0.164) | **0.56** (0.172) |
|  | 0.1 | **0.67** (0.166) | **0.65** (0.170) |
|  | 0.2 | **0.92** (0.261) | **0.81** (0.238) |
|  | 0.3 | **1.22** (0.358) | **1.05** (0.292) |
|  | 0.4 | **1.49** (0.468) | **1.31** (0.400) |
|  | 0.5 | **1.74** (0.490) | **1.49** (0.434) |
| TCGA | 0 | **0.24** (0.056) | **0.25** (0.055) |
|  | 0.1 | **0.45** (0.051) | **0.39** (0.049) |
|  | 0.2 | **0.69** (0.068) | **0.55** (0.053) |
|  | 0.3 | **0.93** (0.089) | **0.72** (0.060) |
|  | 0.4 | **1.17** (0.113) | **0.88** (0.068) |
|  | 0.5 | **1.41** (0.140) | **1.05** (0.077) |
| Yeast | 0 | **0.18** (0.046) | **0.18** (0.046) |
|  | 0.1 | **0.39** (0.045) | **0.32** (0.040) |
|  | 0.2 | **0.63** (0.065) | **0.47** (0.044) |
|  | 0.3 | **0.87** (0.087) | **0.65** (0.056) |
|  | 0.4 | **1.11** (0.113) | **0.81** (0.062) |
|  | 0.5 | **1.35** (0.147) | **0.98** (0.069) |

**Supplementary Figure S3:**
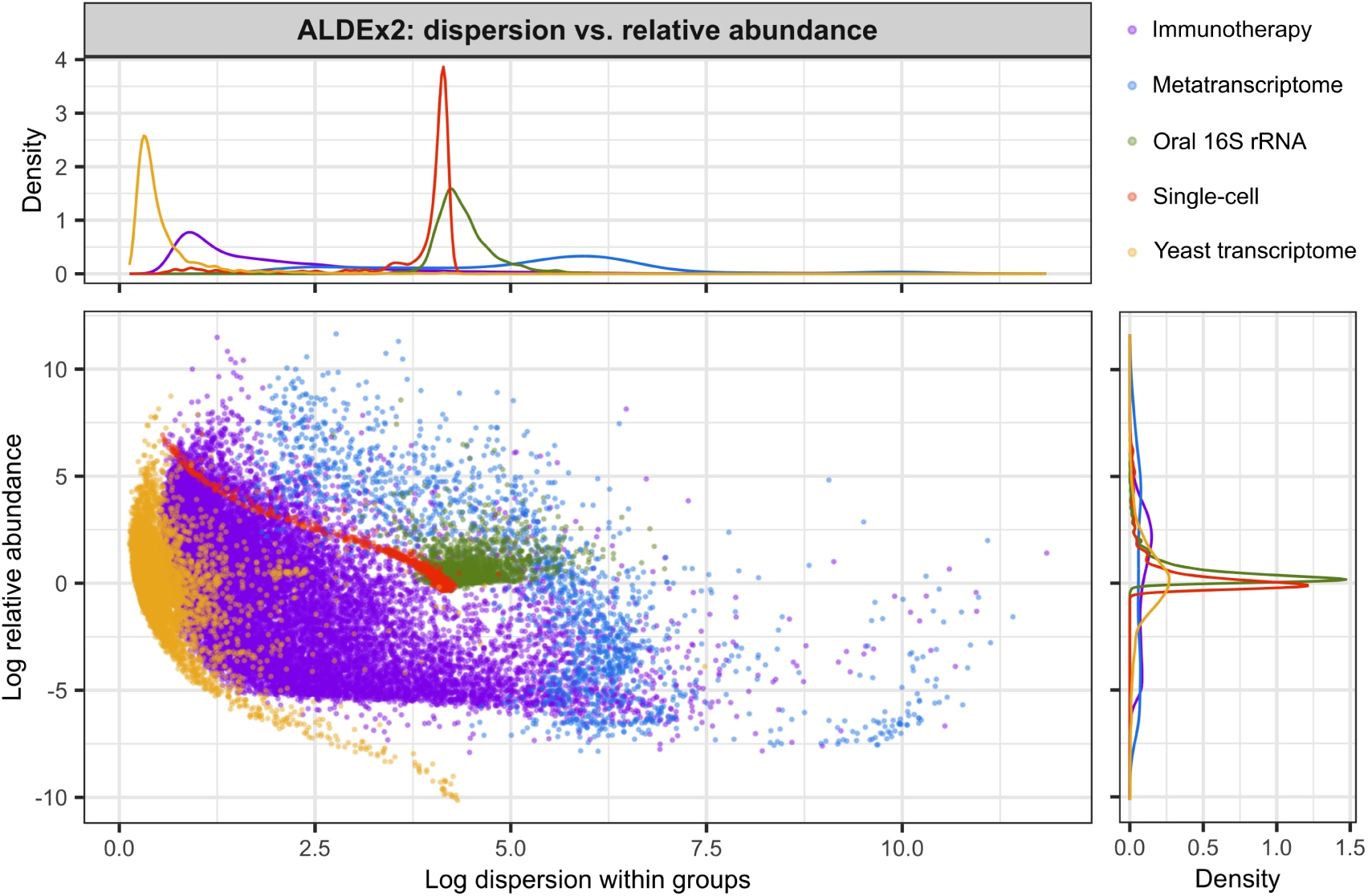
Features from single-cell and metatranscriptome datasets have a larger dispersion compared to bulk RNA-seq data: A single iteration of the ALDEx2 results was selected at random for the single-cell, metatranscriptome, PD1 immunotherapy, yeast transcriptome and oral 16S rRNA datasets and the within-group log dispersion of each feature was plotted against the log abundance.

**Supplementary Figure S4:**
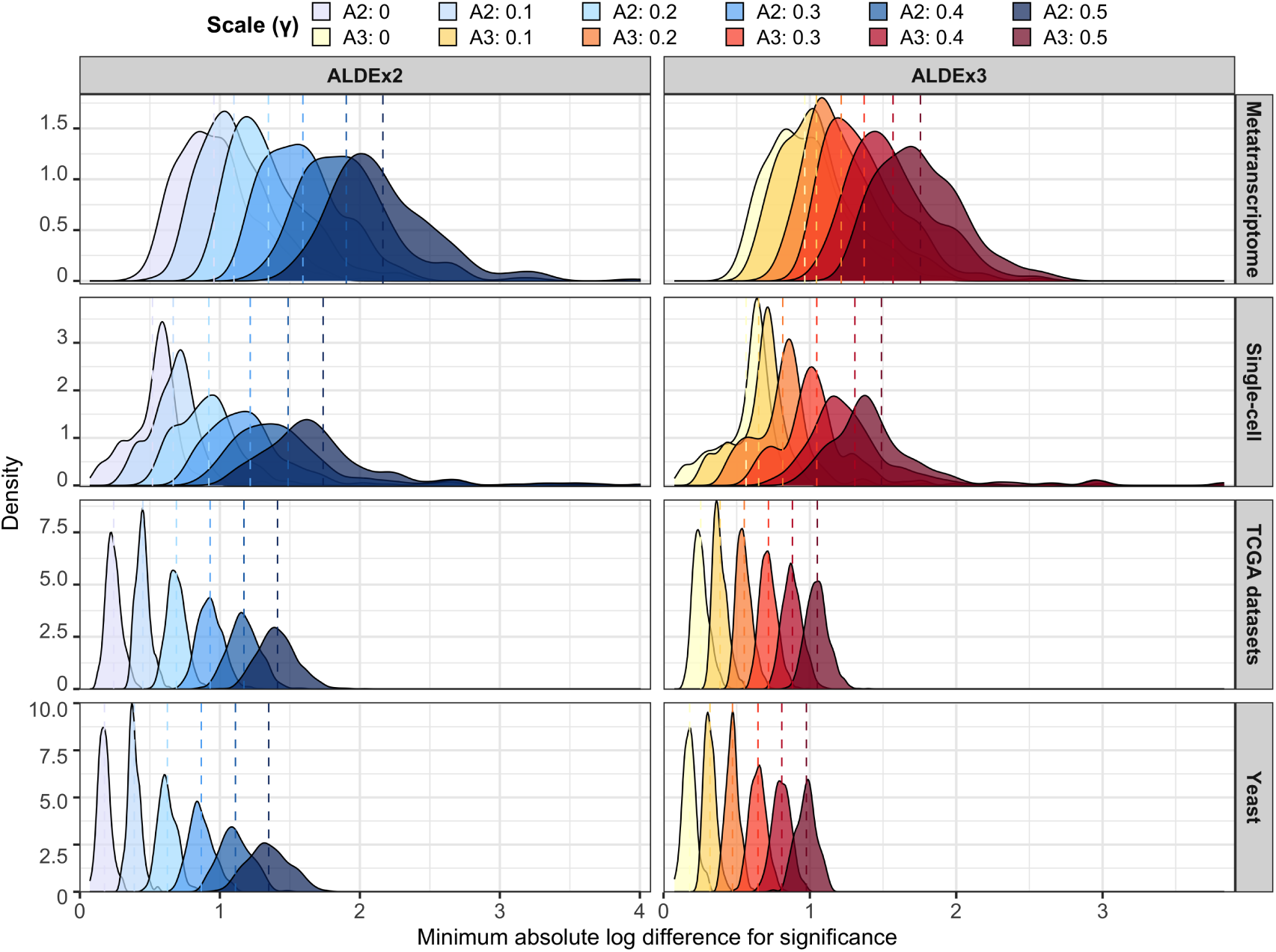
ALDEx3 returns a tighter distribution of minimum log-difference values required for significance across all datasets: For each of the 100 iterations of the given dataset, the minimum absolute difference between groups needed for a gene to be reported as differentially expressed by ALDEx2 and ALDEx3 was recorded, based on a BH-adjusted *P*-value of *<*0.05, for γ values ranging from 0 to 0.5. Data for the Cancer Genome Atlas (TCGA), metatranscriptome, single-cell, and yeast datasets are expressed as density plots and coloured by tool and γ value. Values for TCGA datasets represent an aggregate across the individual tumour transcriptome datasets. Dashed lines indicate the mean value.

**Supplementary Table S2:**
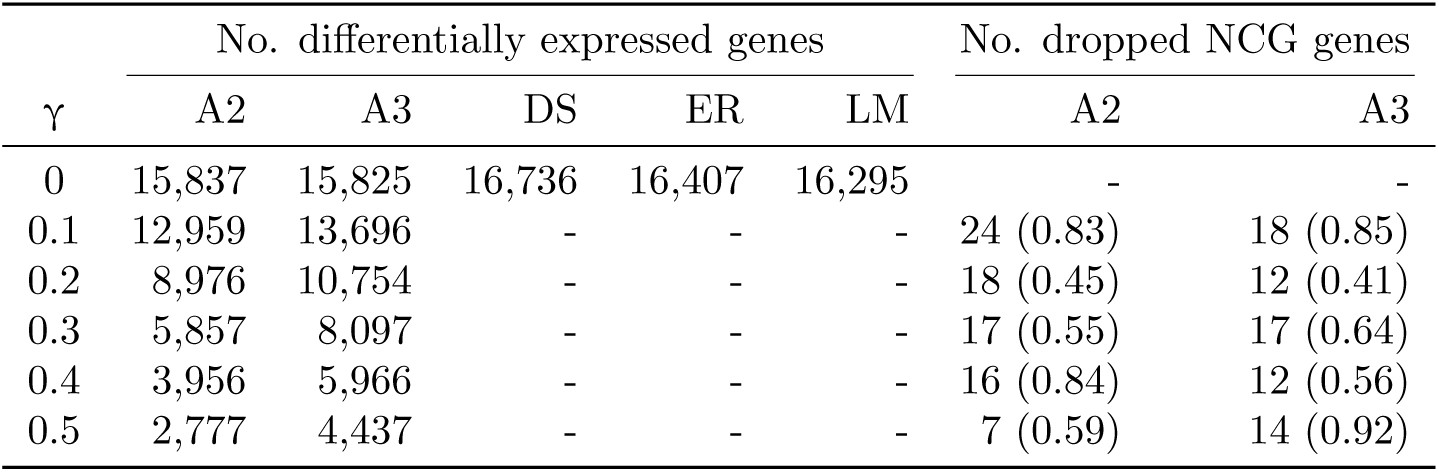
Differentially expressed genes reported by all tested tools for the original, unpermuted BRCA dataset. The total number of genes identified as differentially expressed by ALDEx2 (A2), ALDEx3 (A3), DESeq2 (DS), edgeR (ER) and limma-voom (LM) are shown for increasing values of γ. At each γ value from 0.1 to 0.5, the number and proportion of genes belonging to the ‘canonical drivers’ and ‘breast cancer’ categories of the Network of Cancer Genes [61] that are no longer significantly different between groups (i.e. dropped genes) are also shown for ALDEx2 and ALDEx3. Data represent the results of a single analysis performed with the original, unpermuted count table and group membership vector. Dashes indicate non-applicable fields.

| $\gamma$ | No. differentially expressed genes | | | | | No. dropped NCG genes | |
| --- | --- | --- | --- | --- | --- | --- | --- |
|  | A2 | A3 | DS | ER | LM | A2 | A3 |
| 0 | 15,837 | 15,825 | 16,736 | 16,407 | 16,295 | - | - |
| 0.1 | 12,959 | 13,696 | - | - | - | 24 (0.83) | 18 (0.85) |
| 0.2 | 8,976 | 10,754 | - | - | - | 18 (0.45) | 12 (0.41) |
| 0.3 | 5,857 | 8,097 | - | - | - | 17 (0.55) | 17 (0.64) |
| 0.4 | 3,956 | 5,966 | - | - | - | 16 (0.84) | 12 (0.56) |
| 0.5 | 2,777 | 4,437 | - | - | - | 7 (0.59) | 14 (0.92) |

**Supplementary Figure S5:**
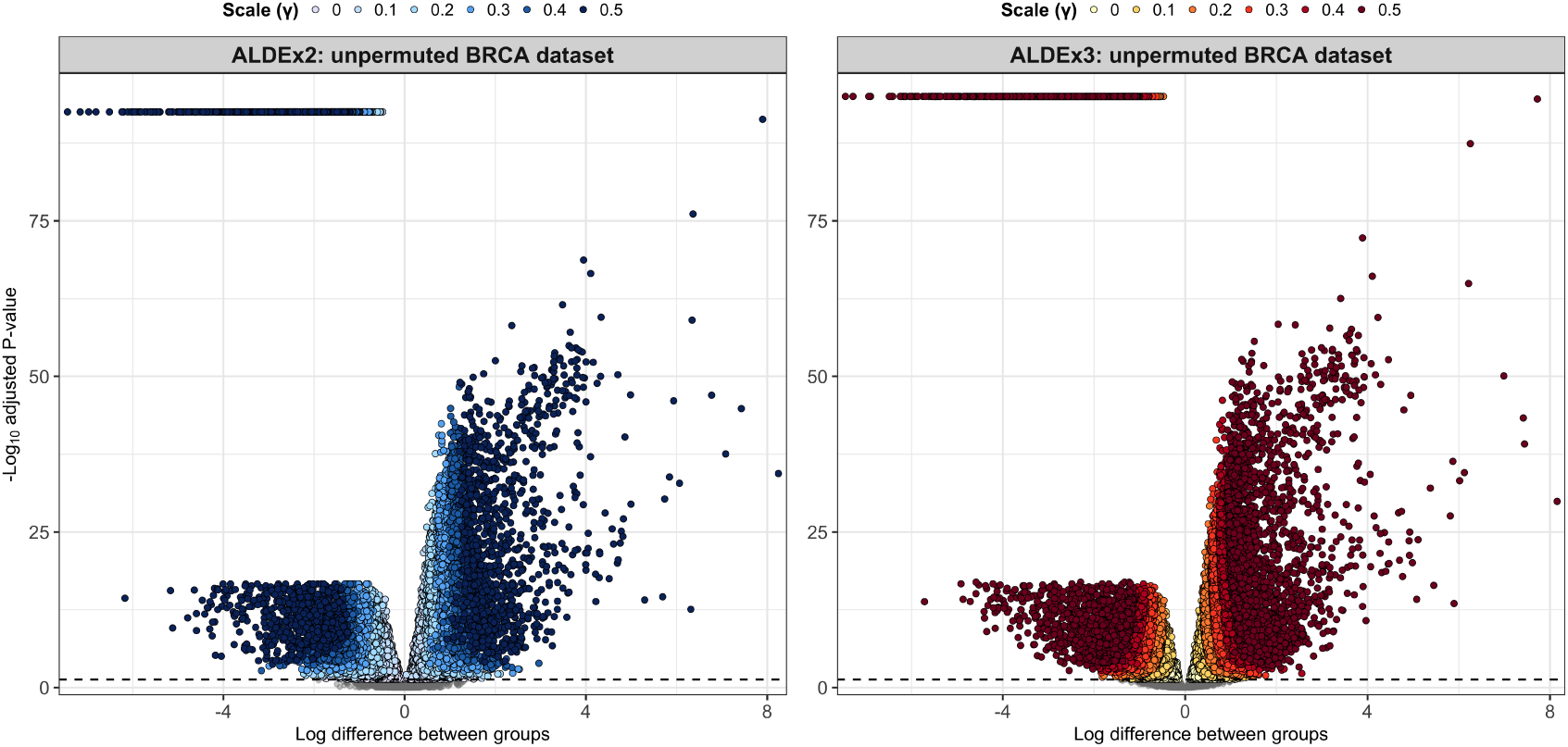
Boundaries between features reported as differentially expressed at different γ values are curved: Volcano plots of log difference between groups vs. −log_10_ *p*-values were generated for ALDEx2 (**left**) and ALDEx3 (**right**) analyses of the original, unpermuted BRCA dataset from the Cancer Genome Atlas. Curved boundaries between points identified as differentially expressed at different γ values can be clearly seen. Each data point represents a feature from the scale-naïve analysis (γ = 0.001). Grey points indicate non-significant features; coloured points indicate features which are significantly different after a certain amount of scale uncertainty has been added to the model. Grey dashed lines indicate Benjamini-Hochberg-corrected *P*-value *<*0.05.

**Supplementary Figure S6:**
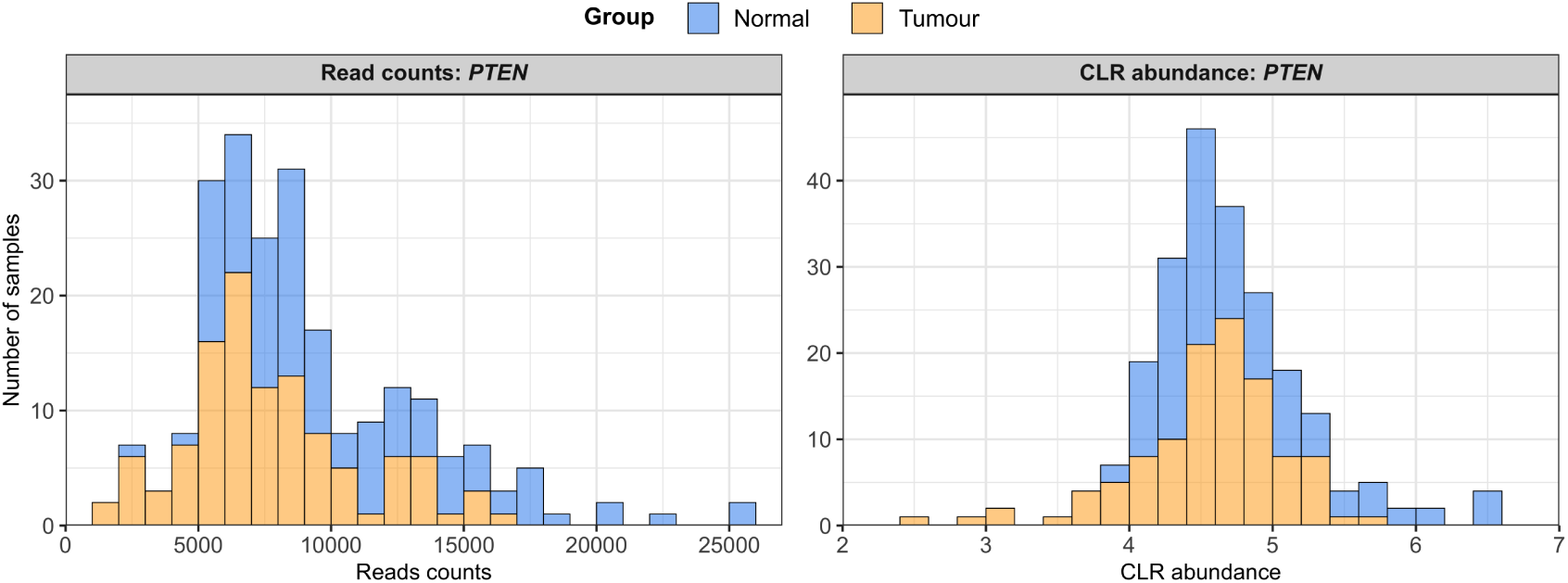
Significant differences in *PTEN* expression are driven by a few outlier samples: Expression of the tumour suppressor gene, *PTEN*, in the Cancer Genome Atlas BRCA dataset is shown in histograms of read counts (**left**) and centred log-ratio (CLR) abundance values calculated by ALDEx2 (**right**). Bars are coloured by sample group. Bin width is 1,000 (reads) or 0.2 (CLR). The limited number of outlier samples in each group can clearly be seen at either end of the histograms, with most samples overlapping in the centre.

## 7 Acknowledgments

We would like to express our thanks to the original authors of all the datasets used in this study for making their data freely and easily accessible for others to use. Their commitment to open science is laudable and is very much appreciated. Specific thanks to the Cancer Genome Atlas research consortium and the donors who contributed samples to their work.

